# Worldwide genetic variation of the IGHV and TRBV immune receptor gene families in humans

**DOI:** 10.1101/155440

**Authors:** Shishi Luo, Jane A. Yu, Heng Li, Yun S. Song

## Abstract

The immunoglobulin heavy variable (IGHV) and T cell beta variable (TRBV) loci are among the most complex and variable regions in the human genome. Generated through a process of gene duplication/deletion and diversification, these loci can vary extensively between individuals in copy number and contain genes that are highly similar, making their analysis technically challenging. Here, we present a comprehensive study of the functional gene segments in the IGHV and TRBV loci, quantifying their copy number and single nucleotide variation in a globally diverse sample of 109 (IGHV) and 286 (TRBV) humans from over a hundred populations. We find that the IGHV and TRBV gene families exhibit starkly different patterns of variation. In particular, with hundreds of copy number haplotypes (instances that have differences in the number of functional gene segments), the IGHV locus has undergone more frequent gene duplication/deletion compared to the TRBV locus, which has only a few copy number haplotypes. In contrast, the TRBV locus has a greater or at least equal propensity to mutate, as evidenced by greater single nucleotide variation, compared to the IGHV locus. Thus, despite common molecular and functional characteristics, the genes that comprise the IGHV and TRBV loci have evolved in strikingly different ways. As well as providing insight into the different evolutionary paths the IGHV and TRBV loci have taken, our results are also important to the adaptive immune repertoire sequencing community, where the lack of frequencies of common alleles and copy number variants is hampering existing analytical pipelines.

## Introduction

By some estimates, genomic variation due to copy number differences underlies more variation in the human genome than that due to single nucleotide differences (1). Yet copy number variation remains challenging to quantify and analyze. Nowhere is this more true than in genomic regions that contain gene families: collections of genes formed through the process of duplication/deletion and diversification of contiguous stretches of DNA (2). Two gene families that are of particular biomedical relevance but for which variation is not well characterized are the immunoglobulin heavy variable family (IGHV), a 1 Mb locus located on chromosome 14 (3, 4), and the T cell receptor beta variable family (TRBV), a 500 kb locus located on chromosome 7 (5). Both loci undergo VDJ recombination, providing the V (variable) component in the biosynthesis of adaptive immune receptors: the IGHV for the heavy chain of the B cell receptor and the TRBV for the beta chain of the T cell receptor (6). In the human genome, both loci are organized as a series of approximately 45 functional V gene segments and are adjacent to a collection of D (diversity) and J (joining) segments. Both loci are present in the genomes of all vertebrates known to have an adaptive immune system, although the arrangement of the IGHV locus can differ between species (7–9). Indeed, the genes comprising the IGHV and TRBV loci are distant paralogs, and are believed to derive from a common ancestral locus in a vertebrate contemporaneous with or predating jawed fishes. That these two loci share genomic features and evolutionary origins makes them an ideal system for a comparative study in gene family evolution.

Here we present the largest investigation to date of genetic variation in the IGHV and TRBV loci using short-read sequencing data. We apply a customized genotyping pipeline (based on (10)) to data from the Simons Genome Diversity Project (SGDP) (11), which performed whole-genome sequencing of a globally diverse sample of human individuals from over a hundred populations. Such characterization of genetic variation in the immune receptor loci sheds light on how the two loci evolved from their common origins. Quantification of variation is also needed in the burgeoning field of computational immunology (12, 13), where the relative abundances of germline variants will help in determining clonal lineages from VDJ sequences. The common copy number polymorphisms we find in our data agree with what has previously been documented, and the most frequent allele we report for each gene segment corresponds to the first or second allele recorded for the gene segment.

## Results

A brief note about gene nomenclature: for the bioinformatic analysis, it was necessary to group together gene segments that are operationally indistinguishable but which have distinct names because they occupy physically different positions on the genome. Our departure from this standard nomenclature is detailed in the Materials and Methods and is also explained where needed below.

To minimize confusion around terminology, we use *polymorphism* as a general term for a genomic unit (nucleotide position or gene segment) that exhibits variation between genomes. Different instances of a particular polymorphism are called *variants*, e.g. a single nucleotide variant or a gene copy number variant. The term *allele* is reserved exclusively for referring to variants of a gene, as in the allele IGHV1-69*01, which is a gene-length variant of the IGHV1-69 gene segment and which may differ in more than a single nucleotide from other alleles of IGHV1-69. This is in line with the usage in the immunogenetics community. We use *haplotype* to refer to the set of operationally distinguishable gene segments that are inherited from a single parent. For example, an individual has two haplotypes of the IGHV locus with one haplotype carrying, say, one copy of IGHV1-69 and the other carrying two copies, say IGHV1-69 and IGHV1-69D.

### General patterns of variation suggest distinct evolutionary dynamics

The evolutionary process of gene duplication/deletion appears to have been faster in the IGHV locus than in the TRBV locus. This is evident in the greater variation in the number of operationally distinguishable IGHV gene segments than in TRBV gene segments (Figure 1A, SI Appendix, Fig. S1). In contrast, the process of diversification — as measured by the number of single nucleotide polymorphisms (SNPs) per gene segment — seems to be comparable or possibly faster in the TRBV locus than in the IGHV locus (Figure 1B). To be conservative, this single nucleotide polymorphism analysis was also restricted to segments which are predominantly two copies per individual and which do not share subsequences with other segments. Our finding that the TRBV locus undergoes less frequent rapid gene duplication/deletions and more frequent nucleotide mutations than the IGHV locus may therefore hold for other vertebrates with adaptive immune systems.

**Figure 1.**
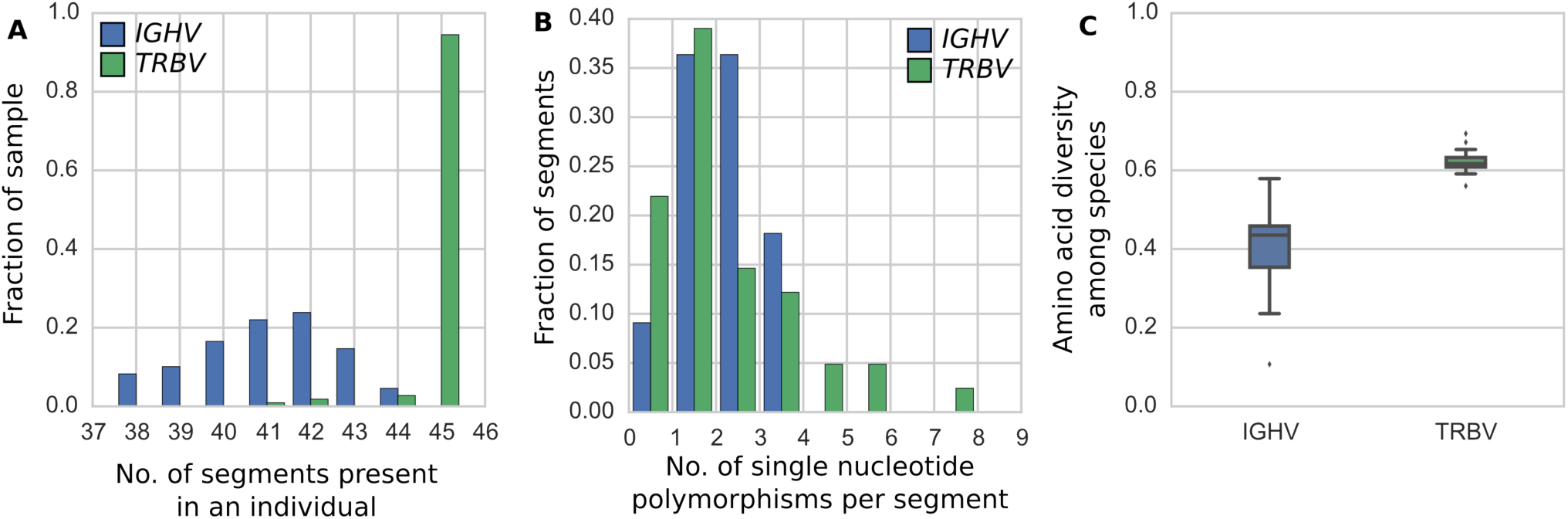
IGHV and TRBV segments exhibit different patterns of copy number and single nucleotide variation, consistent with the genetic diversity in these gene families observed across species. (A) The number of distinct IGHV (blue) and TRBV (green) segments present in each of the 109 individuals from blood and saliva samples. SI Appendix, Fig. S1 shows (A) for TRBV in the full set of 286 individuals. (B) The number of single nucleotide polymorphisms per two-copy IGHV (blue) and TRBV (green) segment. We report a single nucleotide polymorphism only if it is found in at least two out of the 109 genomes from blood and saliva samples. The two distributions are not statistically significantly different (p value of two-sample Kolmogorov-Smirnov test between the blue and green distribution is 0.97). (C) Box plots for average pairwise diversity between IGHV segments and TRBV segments in each of thirteen vertebrate species. For each species, pairwise alignments of all pairs of IGHV segments and all pairs of TRBV segments were performed using ssw **(22)**, an implementation of the Smith-Waterman algorithm **(23)**.

With these different evolutionary dynamics, we expect the IGHV gene family to comprise genes that are more similar to each other on average than the TRBV gene family. Indeed, this is precisely the pattern we see in the human IGHV and TRBV gene families and across twelve other vertebrate species, including four primates, six non-primate mammals, one reptile, and one fish (Figure 1C, Materials and Methods). We also observe less homology between species for the IGHV gene family compared to the TRBV family (SI Appendix, Fig. S2). This is consistent with our finding that gene duplication and deletion occurs more frequently in the IGHV locus: IGHV homologs that are shared between species are erased more frequently than TRBV homologs.

### Copy number variation

Figure 2 and 3 give frequencies of common copy number polymorphisms of IGHV and TRBV genes, respectively. This, and the more detailed SI Appendix, Figs. S3 and S4, will serve as a useful reference for the computational immunology community.

**Figure 2.**
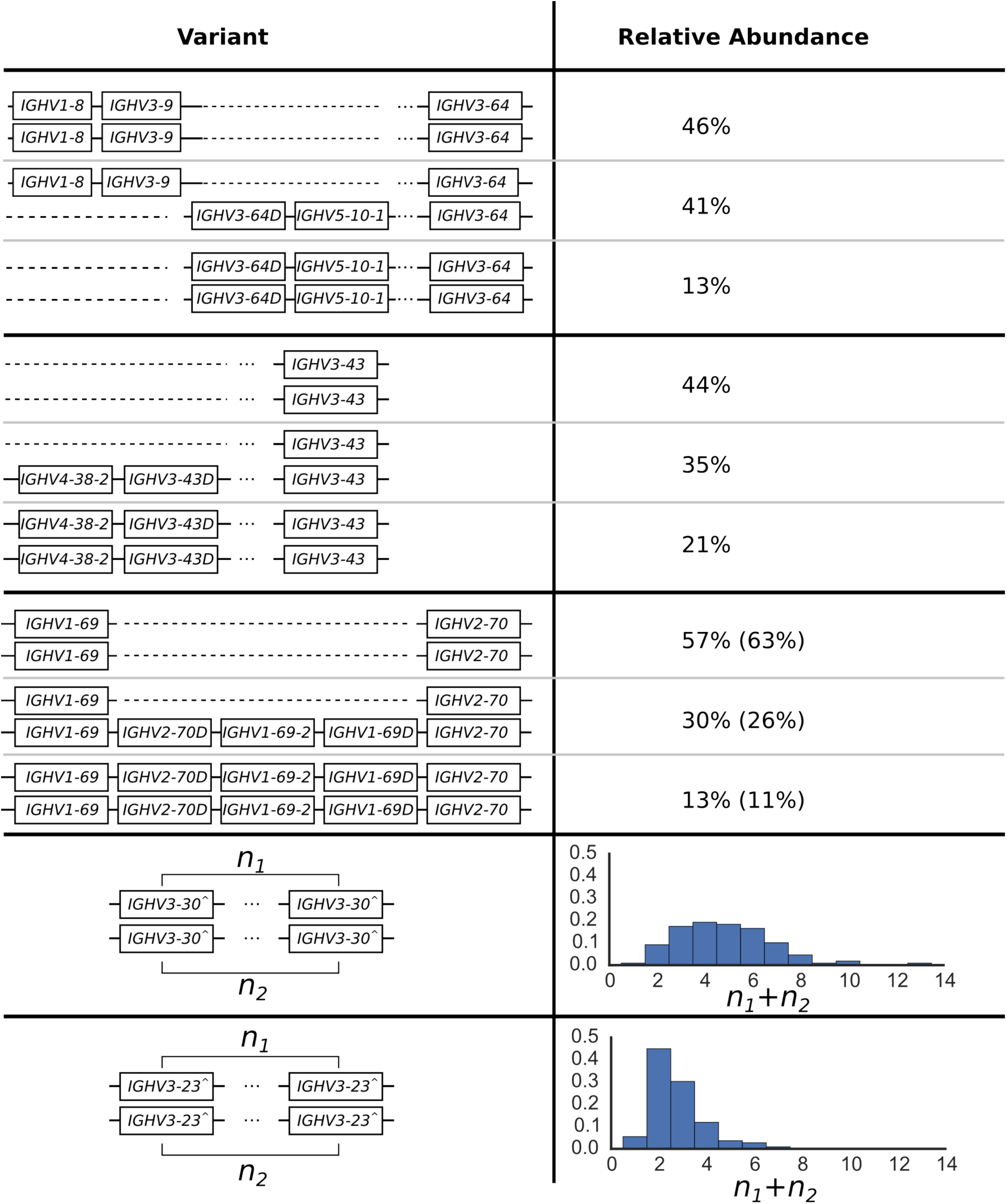
The distribution of common IGHV copy number polymorphisms. Schematics in the left column show the genomic configuration of these polymorphisms determined by previous studies (4). The right column displays the relative abundance in the sample of 109 individuals. For the polymorphism involving IGHV1-69, we also show the relative abundances in the full sample of 286 individuals in parentheses. This is because the impact of VDJ recombination on cell line samples in this J-distal part of the locus has a negligible effect on our copy number estimates. Note that we use IGHV3-30^ as shorthand for the set {IGHV3-30, IGHV3-30-3, IGHV3-30-5, IGHV3-33} and IGHV3-23^ for {IGHV3-23, IGHV3-23D}. The relative abundances for the copy number variants of IGHV3-30^ and IGHV3-23^ are for the total number summed over the two haplotypes in an individual. We estimate the largest haploid number to be 7 for IGHV3-30^ (because the smallest two numbers to sum to 13, the largest copy number called, are 6 and 7) and 4 for IGHV3-23^ (likewise, 4 and 3 are the smallest two numbers to sum to 7).

**Figure 3.**
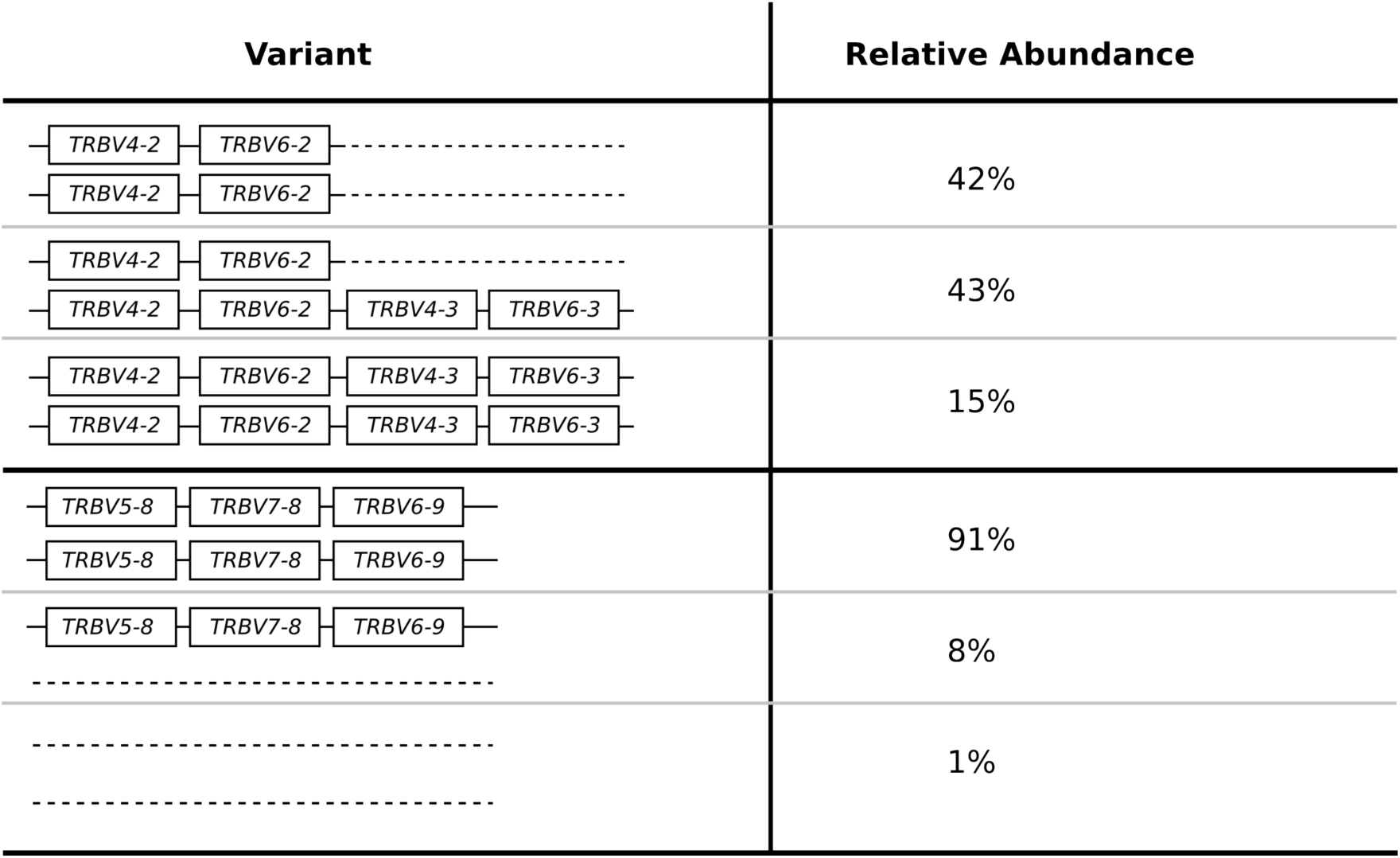
The distribution of common TRBV copy number polymorphisms. Schematics in the left column show putative genomic configurations of the polymorphisms. The right column displays the relative abundance in the full sample of 286 individuals.

#### Lack of geographical associations

In the majority of cases, we find that the distribution of copy number variants within a geographic region is consistent with the global distribution (SI Appendix, Figs. S5 and S6). The two exceptions are: (i) the polymorphism involving IGHV1-69, where the duplication/insertion variant is the major variant among individuals from Africa, despite being a minor variant (28%) of the global sample, and (ii) the three-gene deletion of TRBV5-8, TRBV7-8, and TRBV6-9, which is the major variant among Native Americans, but appears in only 5% of our sample globally. In neither of these two cases is there evidence to suggest the absence of any particular gene is fatal.

#### No correlation between copy number polymorphisms

We find effectively no correlation between copy number polymorphisms in either IGHV or TRBV (SI Appendix, Figs. S7 and S8). The average value of R^2^, the square of the Pearson correlation coefficient, between segments in the different polymorphisms shown in Figure 2 and Figure 3 is 0.021 for the IGHV gene segments and 0.004 for the TRBV gene segments. Thus, the polymorphisms are essentially independent, and we can estimate the number of copy number haplotypes in the two loci. From Figure 2, with three polymorphisms each with 2 (haploid) variants, and with the set {IGHV3-30, IGHV3-30-3, IGHV3-30-5, IGHV3-33} and {IGHV3-23, IGHV3-23D} exhibiting an estimated 7 and 4 (haploid) copy number variants, respectively, this gives approximately 200 IGHV haplotypes (2 x 2 x 2 x 7 x 4), assuming independence between the common copy number polymorphisms. The analogous calculation from Figure 3 for TRBV leads to only a handful of haplotypes (2 x 2).

### Single nucleotide and allelic variation

Because copy number variation in the IGHV and TRBV loci has not been well characterized until now, it has not been straightforward to compare the single nucleotide and allelic variation in gene segments between the two loci. A gene segment with higher copy number could be perceived as exhibiting greater single nucleotide or allelic variation, even though it experiences the same rate of per-base substitution. This is because a duplicate with mutations is difficult to distinguish from a true allele. Having quantified copy number variation of genes across the two loci, we can perform this comparison while minimizing the confounding factor of copy number variation. Specifically, we compare single nucleotide and allelic variation in IGHV and TRBV gene segments that are in two copies in the vast majority of individuals in our sample and for which there is minimal read-mapping ambiguity (11 such IGHV segments, 41 TRBV segments, see SI Appendix, Figs. S3 and S4). We will refer to such gene segments as “two-copy” for short. Several segments in TRBV and in IGHV occur as multi-copies, and some segments have tri- or quadra-allelic SNPs. Our pipeline does not handle calling alleles or single nucleotide variants for these more complex cases.

#### IGHV and TRBV have comparable levels of nucleotide diversity

We find that when restricted to the set of two-copy gene segments in IGHV and TRBV, the two loci have comparable summary measures of single nucleotide and allelic variation (Table 1, Figure 1B). If anything, the TRBV gene segments exhibit greater single nucleotide and allelic diversity. This is seemingly surprising, because if allelic diversity is estimated by taking the average number of alleles per gene segment as per the IMGT database (10), without regard to the segment’s copy number, operationally distinguishable IGHV gene segments have an average of 5 alleles while TRBV gene segments have an average of 2 alleles. The discrepancy in the two ways of estimating allelic variation does not seem to be due to any under-representation of TRBV alleles in the IMGT database: the fraction of putative novel alleles called in our sample is similar between the IGHV and TRBV gene segments (Table 1, third row). As a reference for the antibody repertoire sequencing community, we have provided the relative abundances of alleles for the two-copy gene segments calculated from our sample in SI Appendix, Tables S1 and S2.

**Table 1.** Summary statistics for single nucleotide and allelic variation in IGHV and TRBV. The results tabulated are computed using the same set of 109 individuals and are restricted to the two-copy segments described in the text.

|  | IGHV | TRBV |
| --- | --- | --- |
| Average bp difference per pairs of alleles | 4.6% | 5.1% |
| Average # of SNPs per segment | 1.8 | 2.1 |
| Fraction of novel alleles out of all observed alleles | 45% | 45% |

#### Putatively novel alleles

In addition to restricting our single nucleotide and allelic analysis to two-copy gene segments, we were also conservative in how we called these variants: an allele or single nucleotide variant is called only if it is present in two or more individuals. With this criteria, we called 29 IGHV alleles, of which 13 are putatively novel and 98 TRBV alleles, of which 41 are putatively novel. Of these novel alleles, it is notable that 5 IGHV alleles and 13 TRBV alleles appeared at least 10 times in our sample (we count homozygous alleles as appearing twice; SI Appendix, Table S3). Some of these alleles like IGHV1-45*02_g123A, TRBV10-1*02_g234T, and TRBV12-5*01_c27G are present in high frequency across all geographic regions (putatively novel alleles are named according to the closest matching allele in the IMGT database, followed by point mutations relative to the closest allele). That these novel variants are comprehensively present is further evidence that existing databases of IGHV and TRBV alleles are not yet complete.

#### Single nucleotide/Allelic variants private to geographic regions

We found 5 single nucleotide variants in IGHV gene segments that are private to a single geographic region, and 26 such variants in TRBV gene segments (SI Appendix, Table S4). These variants are not rare: a majority of them are present at greater than 10% frequency in geographic region they are private to, in one case the frequency is 42%. For both loci, the geographic region of Africa had a disproportionate share of such variants: of the five IGHV single nucleotide variants that were private to a geographic region, all five were private to Africa, and of the 26 single nucleotide variants exclusive to a region for TRBV, 20 (77%) were private to Africa. This particular feature of samples from the Africa region is also apparent in our allelic variation analysis (SI Appendix, Table S5). Of the 29 IGHV alleles we called, 5 out of 5 private alleles were private to Africa. Similarly, of the 98 TRBV alleles we called, 15 out of 21 (71.4%) private alleles were private to Africa. These findings of higher levels of diversity primarily in Africa are consistent with prior studies (14) and with the percentage of exon-located single nucleotide polymorphisms that are private to Africa across the entire genome (72.1%). For a complete table of alleles and single nucleotide variants private to a particular region, see SI Appendix, Tables S4 and S5.

#### Geographical clustering of TRBV allelic haplotypes

To investigate whether genetic variation at immune receptor loci exhibit geographical structure, we applied multidimensional scaling to the reconstructed TRBV allelic haplotypes from 286 individuals. Figure 4 illustrates our result, where each point corresponds to an individual. The geographic regions in the legend are those defined by SDGP. SI Appendix, Figs. S9 – S12 show results from applying multidimensional scaling to individuals just from pairs of geographic regions.

**Figure 4.**
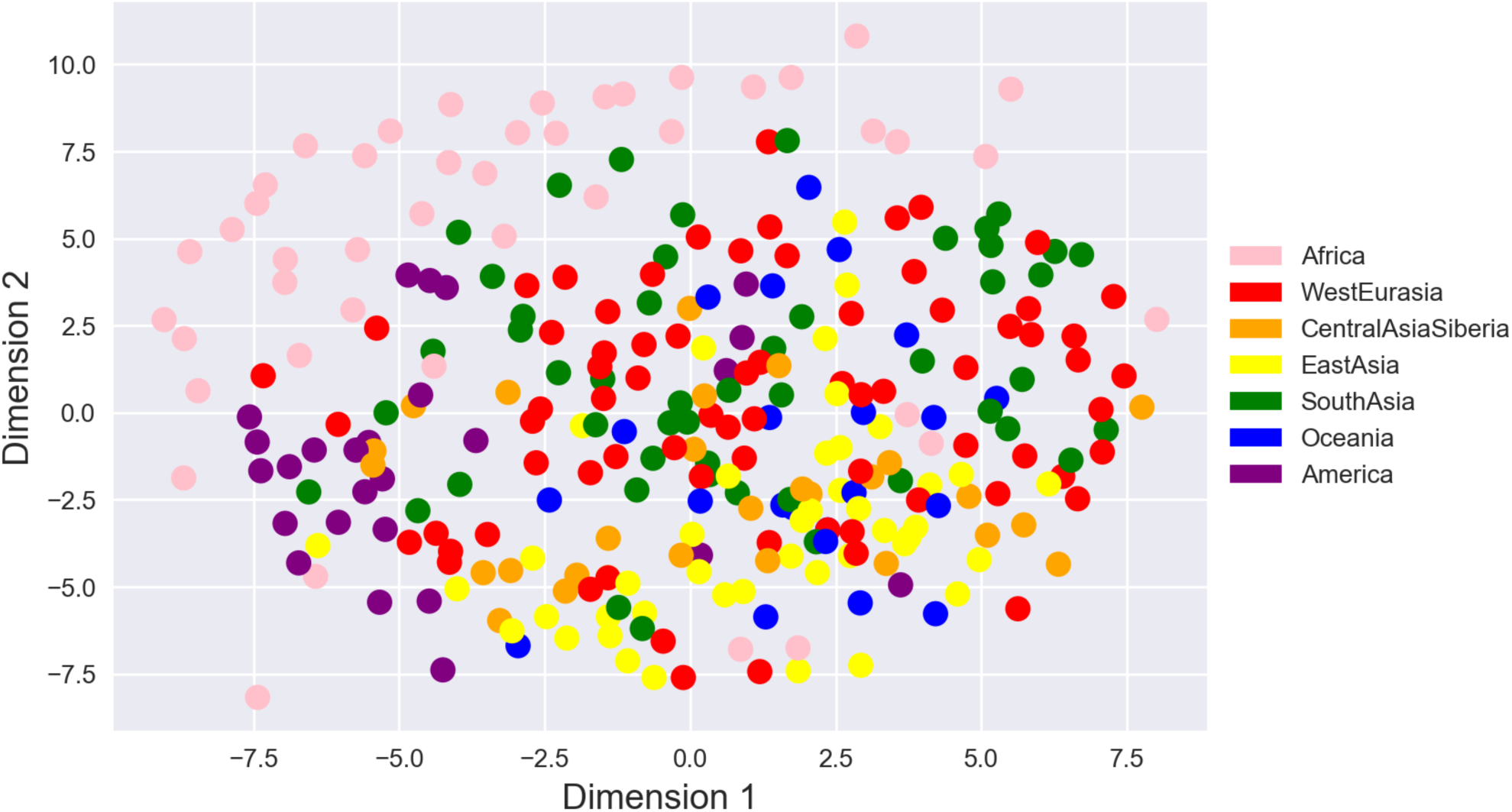
Multidimensional scaling of 286 individuals from all populations (including all DNA source types) in the SGDP dataset, based on their TRBV alleles. The distance between individuals *i* and *j* is defined as the Euclidean distance between vectors *x*_*i*_ and *x*_*j*_, where the *m*th entry in vector *x*_*i*_ is the copy number of allele *m* in individual *i*, taking possible values 0,1, or 2.

As shown in Figure 4, we observed clearest separation between the African population and the rest of the populations, a trend that is also apparent in the pairwise plots (SI Appendix, Fig. S9) and is related to our aforementioned finding that African individuals tend to have the most alleles private to one region. There are a few African individuals who are exceptions to this pattern. Specifically, Masai-1 from Kenya and Saharawi-2 from Morocco consistently cluster more closely with Eurasians (SI Appendix, Fig. S9).

While distinction amongst the other populations is not immediately obvious from Figure 4, every pairwise comparison with Native Americans showed reasonably clear separation from the other populations (SI Appendix, Fig. S10). Additionally, the individuals from Central Asia - Siberia and South Asia (SI Appendix, Fig. S11A) were fairly separable, although the degree of distinction is less prominent compared with those discussed above. Comparisons demonstrating significant overlap include Central Asia – Siberia versus East Asia (SI Appendix, Fig. S11B), West Eurasia versus Central Asia – Siberia (SI Appendix, Fig. S12A), and West Eurasia versus South Asia (SI Appendix, Fig. S12B).

## Discussion

The genetic analysis of gene families remains a technically challenging task in modern genetics. Here, we have made major inroads in calling variants in gene segments in the IGHV and TRBV gene families. We have uncovered patterns of variation that hint at the evolution of the two gene families as well as allelic variants that may be associated with disease specific to a geographic region. Our analysis suggests that the IGHV gene family has experienced more frequent gene duplication/deletion relative to the TRBV gene family over macro-evolutionary time scales. The lack of geographical associations for the majority of common copy number polymorphisms in our sample further indicates that IGHV and TRBV copy number variation was established early in the history of *homo sapiens*, and that it is unlikely that the presence of particular IGHV or TRBV gene segments is vital against any region-specific pathogens. In contrast, we found a number of alleles in both gene families to be private to a particular region and at non-trivial frequencies. Such allelic variants may be promising candidates for investigating genetic variants that are beneficial against infectious diseases endemic in a geographic region.

Our analysis of copy number variation has practical implications for germline IGHV haplotyping: approaches for cataloging variation by sequencing the 1 Mb locus in full (4, 15) will need to consider the possibility that even in a sample of hundreds of individuals, there will be copy number differences between a substantial fraction of haplotypes. Indeed, we find that when we draw two individuals at random from our sample, there is a 98% chance that they will have different sets of IGHV segments present or absent, but only an 11% chance they will have different sets of TRBV segments present or absent. This calculation is based on a coarser, more robust measure of copy number haplotype, where we identify each individual by the presence or absence of a segment, and is therefore conservative. These numbers remain approximately the same even when we restrict our comparisons to individuals within geographical regions, again indicating that the presence or absence of functional segments does not segregate by geographic region (Table S6). These results provide quantitative support for the conjecture made in (16) that “no chromosomes contain the same set of V_H_ gene segments,” where V_H_ refers to IGHV.

Our results are also of immediate relevance to the adaptive immune receptor repertoire sequencing community. The greater complexity in the IGHV locus suggests that using data analysis methods interchangeably between T cell receptor sequences and B cell receptor sequences may not be optimal. The majority of TRBV genes are operationally distinguishable and appear as a single copy per haplotype. Since T cell receptors do not undergo further somatic hypermutation, it makes sense to construct so-called “public” T cell receptor repertoires and analyze individual repertoires in relation to common public repertoires. In contrast, the majority of IGHV genes either vary in copy number or share long subsequences in common with other genes/pseudogenes/orphon genes in the IGHV family (SI Appendix, Fig. S3). Furthermore, immunoglobulins undergo genetic modification via somatic mutation. The analysis of the antibody repertoire may therefore need to be customized to each individual.

Many challenges remain in genotyping complex and variable regions such as IGHV and TRBV. Our approach of using short-read data has a major advantage in being scalable to large sample sizes, allowing population frequencies to be calculated. However, other approaches may be more appropriate if the goal is to genotype a single individual at base pair resolution, rather than a large set of individuals at coarser resolution. Another challenge is measuring the rate of nucleotide substitution in IGHV genes, which requires distinguishing between mutations on paralogous regions from true allelic variation. We have adopted a conservative approach here, restricting our calculation to 11 IGHV genes which we are confident are two-copy. However, these 11 genes may not be representative of all the regions of IGHV that are not subject to copy number variation. An approach which can identify larger tracts of IGHV that are structurally conserved across hundreds of individuals will give a better estimate of the nucleotide substitution rate. Regardless, work in the genetic analysis of the IGHV and TRBV loci will be important to biomedical fields and the growth of personalized medicine.

## Materials and Methods

### Gene nomenclature

The following sets of gene segments were considered operationally indistinguishable (often more than 95% nucleotide similarity) for our bioinformatic analysis: {IGHV3-23, IGHV3-23D}, {IGHV3-30, IGHV3-30-3, IGHV3-30-5, IGHV3-33}, {IGHV3-53, IGHV3-66}, {IGHV3-64, IGHV3-64D}, {IGHV1-69, IGHV1-69D}, {IGHV2-70, IGHV2-70D}, {TRBV4-2, TRBV4-3}, {TRBV6-2, TRBV6-3}, {TRBV12-3 TRBV12-4}.

### SGDP Dataset

Whole genome shotgun sequencing reads were collected in a previous study, the Simons Genome Diversity Project (11). Briefly, 300 genomes from 142 subpopulations were sequenced to a median coverage of 42x, with 100 base pair paired-end sequencing on the Illumina HiSeq2000 sequencers. The reads from 286 of these genomes were mapped to the set of functional alleles (IGHV or TRBV) as per the IMGT database (10). Of the 286 individuals, only those from non-cell line, i.e. blood and saliva DNA sources (109 in total), could be used for IGHV analysis. This is because in these cell lines, which are based on immortalized B cells, the IGHV locus is truncated relative to germline configuration due to VDJ recombination. Details of individual samples can be found in Supplementary Data Table 1 of reference (11). For the TRBV locus, we used the full set of 286 genomes. Note: we only had access to 286 genomes of the 300 genomes: 300 minus the 14 individuals with labels SS60044XX.

### Read mapping

For each individual, we retained reads from two separate procedures: (i) reads that map to the IGHV and TRBV loci on the GRCh37 reference, (ii) reads that map to a list of functional IGHV and TRBV (from the online IMGT database (10)).

Procedure (i) used bwa mem (https://github.com/lh3/bwa) with default parameters, e.g.:

~~~
bwa mem grch37.fa read1.fq read2.fq | gzip −3 > aln-pe.sam.gz
~~~

Procedure (ii) used minimap (https://github.com/lh3/minimap):

~~~
minimap -w1 -f1e-9 imgt_ighv.fa.gz read-se.fa.gz > out_ighv.mini
minimap -w1 -f1e-9 imgt_trbv.fa.gz read-se.fa.gz > out_trbv.mini
~~~

Although procedure (i) was sufficient in our previous study (17) for identifying all the IGHV genes in an individual, it was clear that for the Simons dataset, the mapped reads were biased to the GRCh37 reference (SI Appendix, Fig. S13). This is possibly due to differences in mapping algorithms in the different sample sets. For this reason, procedure (ii) was needed to ‘catch’ the reads that are not in the reference genome but which map to known IGHV and TRBV gene segments. Such gene segments include IGHV1-69-2 and TRBV5-8.

### Read filtering

The disadvantage of procedure (ii) is that reads from highly similar pseudogenes and orphon genes may get mixed with reads from functional genes (SI Appendix, Fig. S14). Thus, for each of the IGHV and TRBV loci, we filter the set of raw reads, aiming to minimize reads that have been erroneously mapped to a functional gene segment. This required taking into account idiosyncrasies of individual segments, especially their similarity to pseudogenes and orphon genes. Due to space limitations, the full details of the filtering steps are in the SI Appendix, SI Text.

### Copy number calls/contig assembly

After read filtering, we have, for each individual, a set of reads binned by operationally distinguishable segment. We next run the assembler Spades

(18) to construct a contig for each segment to obtain:

1. kmer coverage for the segment in that individual
2. A first estimate of the nucleotide sequence of the gene in that individual

For example, for a fixed individual, the script we execute to assemble the contig for IGHV6-1 is:

spades.py –k 21 –careful –s IGHV6-1.fastq –o contigs/IGHV6-1

The choice of kmer of size 21 is because it was the longest kmer that ensured successful contig construction for our 100 bp reads at around 40 coverage depth. The kmer coverage is then converted to per-base coverage, scaled to account for the trapezoidal shape of the read coverage profile, and then normalized by the individual’s genome-wide coverage to obtain a point estimate for copy number (details of calculation in SI Appendix, SI Text).

### Variant calls for common polymorphisms

To call the copy number variants for common polymorphisms involving multiple genes, we use the point estimates for copy number. The reason we do not round the point estimates to whole integers is that the rounding process may introduce additional error into our variant calls. For each polymorphism, we performed a hierarchical clustering (scipy.cluster.hierarchy) on the set of individuals, representing each individual as a vector comprised of the copy number estimate for each gene in the polymorphism. The results of the clustering are shown in SI Appendix, Figs. S15-S19.

We looked in the IGHV data for evidence of copy number polymorphisms previously documented in (4). For the TRBV data, the polymorphism involving {TRBV4-2, TRBV4-3} and {TRBV6-2, TRBV6-3} has previously been documented. The polymorphism involving TRBV5-8, TRBV7-8, and TRBV6-9 was identified by first clustering using TRBV5-8 copy number estimates alone, and then noticing that such a clustering also induced a clear-cut partition of the copy number estimates for TRBV7-8 and TRBV6-9.

### Allele and single nucleotide variant calls

The contigs and reads for two-copy segments (those that occurred as single copy in the locus in the vast majority of individuals in our dataset and which do not share substantial nucleotide sequence with any other gene segments) were analyzed for allelic and single nucleotide polymorphisms (SNPs) by phasing these segments for each individual. Because the assembly step in the pipeline produces only one contig, we reconstructed the two distinct allelic sequences on each chromosome through additional steps, which are as follows:

1. Mapped the filtered set of reads to the contig constructed via the customized pipeline using Bowtie2 with the parameters *--local --score-min G,20, 30*.
2. The results from Bowtie2 were fed to GATK (19) for variant calling, producing VCF files identifying polymorphic sites, using the HaplotypeCaller with parameters *-ploidy 2 -stand_call_conf 30 -stand_emit_conf 10*.
3. The variants from GATK were then phased using HapCUT2 (20).

a. When few or no reads covered two variant locations, HapCUT2 failed to phase the full segment. To account for this issue, we took all combinations of completely phased blocks as potential allele sequences. For example, consider a sequence of length three, each position having an unphased polymorphic site with A on one chromosome and T on the other. Taking combinations results in a. four pairs of complementary sequences (AAA, TTT), (ATA, TAT), and (AAT, TTA), and (ATT, TAA).
b. With the resulting pairs of phased sequences, we compute the probability of observing each candidate pair of allele sequences (a_1_, a_2_), which is calculated by taking the maximum of P(a_1_|a_2_)*P*(a_2_) and *P(*a_2_|a_1_)P(a_1_). P(a_x_|a_y_) is the fraction of individuals observed to have both allele sequences a_x_ and a_y_ out of all individuals observed to have allele a_y_ and P(a_x_) is defined as the fraction of observations of a_x_ out of all individuals.
c. If all candidate pairs of allele sequences were not observed in any other individual, then the following was selected as the individual’s phased allele sequence pair: the candidate allele sequence a_x_ with the greatest frequency and its complementary allele sequence to a_x_. If all candidate sequences a_x_ for an individual were not observed in the rest of the population, then no allele sequence was reported for that individual for that segment.
4. To be conservative, we kept only the alleles found in at least two different individuals.

After phasing, we quantified the allelic and single nucleotide variation. Alleles were given IMGT names if their sequence exactly matched an allele in the IMGT database. Otherwise, the name of the closest allele was given with an appended suffix for each mutational difference from the closest IMGT allele. For example, allele ‘ IGHV1-18*01_a168G’ denotes an allele whose sequence is that of IMGT allele IGHV1-18*01, but with a ‘ G’ at position 168 rather than an ‘ A’. Alleles were called “novel” if it differed in at least one nucleotide from an existing IMGT allele. To calculate the average base pair difference per pairs of alleles (as seen in Table 1), for each segment we computed the average base pair difference between all pairs of alleles, and then averaged over all segments. To determine which sites were single nucleotide polymorphisms, all allele sequences for each segment were compared, site-by-site. In our analysis, we focus on the sites with two variants.

### Multidimensional scaling analysis

Multidimensional scaling was performed in Python using the manifold.MDS([n_components, metric, n_init,…]) function from the sklearn.manifold module. The data fit by the model the Euclidean distance between x_i_ and x_j_ where the mth entry in vector x_i_ is the copy number of allele m in individual i, taking possible values 0,1, or 2.

### Other species

Amino acid sequences for IGHV segments and TRBV segments for the interspecies comparison were obtained from vgenereportoire.org (21). Species are: *homo sapiens* (human), *pan troglodytes* (chimpanzee), *gorilla gorilla* (gorilla), *pongo abelii* (orangutan), *macaca mulatta* (rhesus macaque), *mus musculus* (mouse), *canis lupus familiaris* (dog), *oryctolagus cuniculus* (rabbit), *orcinus orca* (orca), *monodelphis domestica* (opossum), *ornithorhynchus anatinus* (platypus), *crocodylus porosus* (crocodile), *danio rerio* (zebrafish).

## Acknowledgments

We gratefully acknowledge David Reich for useful discussions and for making short-read data from the Simons Genome Diversity Project available to us. This research is supported in part by an NIH grant R01-GM094402, and a Packard Fellowship for Science and Engineering. YSS is a Chan Zuckerberg Biohub investigator.

